# Quantifying flight aptitude variation in wild *A. gambiae* s.l. in order to identify long-distance migrants

**DOI:** 10.1101/2020.03.03.975243

**Authors:** Roy Faiman, Alpha S. Yaro, Moussa Diallo, Adama Dao, Samake Djibril, Zana L. Sanogo, Margery Sullivan, Asha Krishna, Benjamin J. Krajacich, Tovi Lehmann

## Abstract

**Background:** In the West African Sahel, during the 5-7 month-long dry season mosquito reproduction is halted due to the absence of surface waters required for larval development. Recent studies have suggested that both *Anopheles gambiae* s.s and *A. arabiensis* persist in this region by migration from distant locations where larval sites are perennial, and *A. coluzzii* engages in long-distance migration, presumably within the Sahel, following shifting resources due to the ever-changing patterns of Sahelian rainfall. Understanding mosquito migration is key to malaria control, a disease that still kills >400,000 people, mostly children in Africa.

**Methods:** We used a new tethered-flight assay to characterize flight in the three primary malaria vectors mentioned above and evaluated seasonal differences in their flight activity. The flight of tethered wild mosquitoes was audio-recorded from 21:00h to 05:00h in the following morning and three flight aptitude indices were examined: total flight duration, longest flight bout, and the number of flight bouts during the assay. Based on recent studies, we predicted that *(i)* the distribution of the flight aptitude indices would exhibit bi-modality and/or marked skewness, indicating a subpopulation of high flight activity (HFA) associated with long-distance migrants, in contrast to low flight activity (LFA) in appetitive flyers. Additionally, flight aptitude would *(ii)* increase in the wet season, (*iii*) increase in gravid females, and *(iv)* vary among the vector species.

**Results:** The distributions of all flight indices departed sharply from a normal curve, and were strongly skewed to the right, consistent with the division of the population into a majority of LFAs and a minority of HFAs, e.g., the median total flight was 586 seconds, and its maximum value was 16,110 seconds (~4.5 h). As predicted, flight aptitude peaked in the wet season and was higher in gravid females than in non-bloodfed females. Flight aptitude was higher in *A. coluzzii* than in *A. arabiensis*, but *A. gambiae* s.s. was not statistically different from either. We evaluated differences in wing size and shape between LFAs and HFAs. During the wet season, wing size of HFA *A. coluzzii* was larger than that of LFAs; it was wider than predicted by its length, indicating a shape change. However, no statistically significant differences were found in wings of *A. gambiae* s.s. or *A. arabiensis*.

**Conclusions:** The partial agreement between the assay results and predictions suggest a degree of discrimination between appetitive flyers and long-distance migrants. Wing size and shape seems to indicate higher flight activity in *A. coluzzii* during the wet season.

## Introduction

The long-distance migration (LDM) of insects (Dingle 2014; Dingle and Drake 2007; Chapman, Reynolds and Wilson 2015; Chapman and Drake 2019) produces numerous services and disservices relevant to human agriculture, economy and health (Bauer et al. 2017), e.g. in terms of food security (Rainey 1989; Leskinen et al. 2011; Hu et al. 2019; Glick 1939; Dingle 2014; Cheke et al. 1990; Chapman, Reynolds, and Smith 2004), public health (Garms, Walsh and Davies, 1979; Ritchie & Rochester, 2001; Sellers, 1980) and even the transfer of nutrients by migrating insects (Hu et al. 2016; Landry and Parrott, 2016; Wotton et al. 2019). Here, we follow the definition of migration as persistent movements of individuals that are not affected by immediate cues for food, reproduction, or shelter (i.e. ‘station-keeping’), and which have a probability to take the animal to a new environment suitable for survival/breeding (Gatehouse 1997; Dingle and Drake, 2007a; Chapman, Reynolds and Wilson, 2015). We are particularly concerned with the Sahelian Zone of West Africa, where mass seasonal nocturnal migrations of pest insects such as grasshoppers and pyrrhocorid bugs, into- and back out of the Sahel in Mali and Niger have been described. These follow the seasonal shifts in wind direction as the Inter-Tropical Convergence Zone (ITCZ) moves north and then south (Maiga, Lecoq, and Kooyman 2008; Riley and Reynolds 1983; 1979b; Duviard, 1977), with the migrants taking advantage of ephemeral, but seasonally-available, habitats. However, because it is so much easier to notice immigration into areas depleted of conspecific populations, many other cases of insect migration may have been discounted.

Anecdotal evidence from various regions of the world suggested that mosquitoes too could engage in long-range dispersals on the wind (Reynolds et al. 1999; Reynolds, Chapman and Harrington 2006; Ming et al. 1993; Ming and Xue 1996; Johansen et al. 2003), but the prevailing view held that such movements are accidental in most disease vector species and thus of negligible epidemiological significance (Service 1993; 1997). However, a recent aerial sampling study showed that in the Sahel, many species of mosquitoes, including *An. coluzzii* and *An. gambiae* s.s., as well as several secondary malaria vectors regularly engage in seasonal flights 40-290 m above ground (Huestis et al. 2019). Because of the large number of migrants, most of which were gravid females, and the large distances they were able to cover, the likely epidemiological significance of such movements was inescapable.

As a previously unrecognized behavior in malaria mosquitoes, windborne long-distance migration raises new questions including the following: Are all mosquitoes long-distance migrants or do only a fraction of the population migrate? Are migrants more common in some species and under certain conditions, and if so, what are these? And, what are the physiological and molecular mechanisms involved in preparation for and during the journey? Addressing such questions using aerial sampling would be challenging for many insect species because they will be intercepted at a frequency of less than one per sampling night. Tethered-flight mills (Minter et al. 2018) have been used effectively to characterize short- and long-distance flyers in many species (e.g. cotton strainers (*Dysdercus fasciatus*, Pyrrhocoridae) (Dingle and Arora 1973), corn leafhoppers (*Dalbulus maidis*, Cicadellidae) (Taylor et al. 1992), the brown marmorated stinkbug (*Halyomorpha halys*, Pentatomidae) (Lee and Leskey 2015) and Buprestid beetles (Taylor et al. 2010), facilitating investigations to address questions such as those mentioned above. However, despite their intuitive appeal, flight mill results have also been reported to be at odds with expectations in species with well-established migration (Colvin and Gatehouse 1993; Jones et al. 2016; Dällenbach et al. 2018). Thus, the approach has its value and pitfalls (Taylor et al. 2010; Minter et al. 2018; Naranjo 2019) and predicting when it would be useful is not always clear. Additionally, flight mills are well suited for laboratory studies, but they can be challenging for field experiments. Therefore, we developed a novel assay to measure flight of tethered mosquitoes under field conditions. As has been done using flight mills, flight aptitude measures, such as total time in flight during the assay may help distinguish between persistent flight, which is presumably enhanced in migrants which would fly considerably longer than ‘appetitive’ flyers. Therefore, we sought to estimate the fraction of strong flyers (presumed migrants) among wild mosquitoes, representing different species at different seasons. Based on population dynamics results (Adamou et al. 2011; Dao et al. 2014; Lehmann et al. 2010), we initially predicted a high fraction of migrants during the early and late rainy season and lower fraction of migrants in *A. coluzzii* compared with *A. gambiae s.s.* and *A. arabiensis*. However, recent aerial sampling data (Huestis et al. 2019) has led us to modify the latter prediction, to suggest the presence of migrants across the three species. The data from the aerial sampling study (Huestis et al. 2019) also led us to modify our initial prediction reflecting the high frequency of gravid females in altitude. Finally, we investigated whether putative migrants (based on our flight mill data) exhibit different wing morphology.

## Materials and Methods

### Study area

Tethered-flight assays were conducted in the Sahelian village of Thierola (−7.2147 E, 13.6586 N) from August 21, 2015 until November 21, 2015 and from March 28 until September 27, 2016. During the dry season, when mosquitoes were too scarce in the Thierola area, assays were conducted in the rice-cultivation town of Kangaré (Selingue commune) (−8.198 E, 11.644 N, 250 km SSW of Thierola), between December 24, 2015 and February 12, 2016. A total of 114 assay-nights was conducted in the field, throughout the course of a year: during the rainy season (June-October), and during the dry season spanning November-April. In both villages, flight assays were conducted indoors, within local houses that were selected for experimentation. Windows in the experiment rooms enabled limited natural light but no wind or air currents. In Thierola the mean nightly (21:00 to 5:00h) temperature throughout the rainy season was 24.5°C (range: 20.2-32.4°C) with a mean RH of 88.6% (range: 31-100%). During the dry season the mean nightly temperature was 23.4°C (range: 10.9-35.6°C) with a mean RH of 27.3% (range: 5.0-100%). In Kangaré the mean nightly temperature between December and February was 26.2°C (range: 24.5-27.7°C) with a mean RH of 29.3% (range: 20.9-63.1%).

### Mosquitoes

Wild *A. gambiae* s.l. were collected both indoors and outdoors, within the village, using aspirators, on the morning of a flight assay (between 07:00-10:00h) and were kept indoors in 1-gallon plastic cages covered with a dampened cloth. Mosquitoes were provided water on a cotton ball for hydration until 16:00h. Before each flight assay, active mosquitoes (reacting with flight to tapping on the cage) were selected by gonotrophic stage (gravid or unfed) which was assessed by visual examination of the abdomen.

Following morphological identification (Gillies and De Meillon 1968), only *A. gambiae* (sensu lato) mosquitoes were included in the flight assays. Subsequent species identification was performed by species-specific PCR and PCR-RFLP with legs as template (Fanello, Santolamazza, and Della Torre 2002). Thirteen individuals not identified by this assay were excluded.

### Wing measurements

Wing length (WL) and wing width were measured as described elsewhere (Diana L Huestis et al. 2011). Briefly, wings mounted under a coverslip with glycerol were photographed at x25 magnification using a microscope (Olympus DM-4500B) coupled with a digital camera (MC170 HD, Leica Microsystems, Wetzlar, Germany). For each wing, 14 specific landmarks (Fig. S1), i.e. vein intersections were mapped using the tps-DIG32 2.15 software package (Rohlf, 2010). Wing length was measured between points 1 and 10, Fig. S1). Wing width was the average value of the height of three triangles formed between landmarks: 1-2-13 (proximal triangle), 2-5-11 (medial triangle), and 2-12-14 (distal triangle) (Fig. S1). For ~265 damaged wings, WL was predicted using a regression analysis based on distances between landmarks 1 and 7 or 4 and 10, for wings with damaged tip or base, respectively (see Sup. Meth.). Regression models showed that these predictors accounted for >95% of the variation in wing length based on intact wings.

### Tethered flight assay

Individual mosquitoes were gently aspirated from the cage and transferred into a 1.6 ml microcentrifuge tube with the bottom removed and replaced with muslin netting. This tube was inserted into a 50 ml Falcon tube containing a cotton ball with 2-3 drops of diethyl-ether at the bottom of the tube. Mosquitoes were anesthetized by 3-4 sec exposure to the ether-vapor rich environment and quickly placed, wings down, under a dissection stereo-microscope (Zeiss Stemi 2000-C. Carl Zeiss Microscopy, Germany). Using entomological pins (Morpho No.3. Ento Sphinx, Czech Republic), sharp ends clipped off, pins were bent twice at 90°(to a square bracket shape). The tip end of the pin was lightly dipped in glue (Elmer’s, Glue-All E1322, Atlanta, GA) and gently pressed on to the ventral side of the posterior abdomen (covering the lower half of the abdomen). The other end was threaded through the base of a disposable 10 μl pipette tip of which the nozzle cone (dispersing end) was cut off (Fig. S2a). Tether pins were cut to size and bent enabling all legs to remain suspended in the air throughout the assay, thus preventing tarsal contact and flight cessation. Tethered mosquitoes were allowed 2 hours to fully recuperate from the anesthesia before the flight assay (moving of legs and wings usually began 2-3 minutes after removal from the ether), during which time their fore-legs were allowed to rest on a folded piece of paper (=‘leg-rest’) providing tarsal contact and preventing flight (Fig. S2b).

At the end of the recuperation time, tethered mosquitoes were inserted into individual flight tubes (50 ml Falcon®, Corning, NY, USA) positioned within a polyurethane foam hive (foam mattress) for soundproofing and environmental cue reduction (Fig. S2c). Tether constructs (mosquito, pin and base) were secured onto a small piece of double-coated urethane foam tape (3M^®^, Cat. No. 4026. St. Paul, MN) to fasten them 1 cm inward of the flight tube edge. Each flight tube housed a small microphone (ME-15, Olympus America Inc., Center Valley, PA, USA) attached to a portable voice recorder (VN-5200PC, Olympus America Inc., Center Valley, PA, USA) (Fig. S2c and c_i_) to record flight sound.

### Flight Sound Extraction

Tethered mosquitoes were recorded over a 10-hour period starting at 21:00h. Sound recordings were downloaded and read using Audacity 2.1.2 open-source software (Mazzoni, 2016). Flight bouts (episodes of flight) were identified visually in spectrogram view (Fig. S3a), and questionable flights were confirmed by listening for flight sound on the soundtrack. For each flight bout, start time and duration were manually recorded into a Microsoft Excel® spreadsheet. Since the shortest time frames measured by the software were 1-second long, all flight bouts shorter than one second were inserted into the database as one second long bouts. In total 216 individual mosquito recordings from 47 different assay-nights were extracted manually. Subsequently, we used Raven Pro 1.5 Interactive Sound Analysis Software (The Cornell Lab of Ornithology 2014) to detect and extract flight bouts from the sound files. Raven Pro utilizes a Band Limited Energy Detector (BLED) which estimates the background noise of a signal and uses this to find sections of the signal that exceed a user-specified signal-to-noise ratio (SNR) threshold in a specific frequency band during a specific time (Duan et al. 2013). BLED outputs were verified audibly or visually in spectrogram view to rule out false positive flight bouts (Fig. S3b).

All 8-hour long sound files (approximately 140 megabytes each) were split into four sections before their analysis in Raven Pro 1.5 to allow modification of the sound detector (BLED) for changing background noises throughout the night (filtering out background noise produced by passing vehicles, electricity generators, crickets, farm animals, rain, etc.), and to enable sufficient computer processor memory for the BLED runs. The Raven Pro sound detectors picked up flight bout lengths as short as 0.01 seconds. Flight bouts separated by rest periods <1.45 seconds picked up by BLED (12%) were pooled to a continuous flight bout to ensure consistency with the manually extracted method (see above). The resultant flight duration value was virtually identical to the original values in total-, mean- and max flight but was consistent with the manual data in respect to flight bouts.

Flight bout records produced by Raven Pro were exported as text files (.txt), which were then read into one large sound database (including the manual flight extraction file) using R-Studio (see database in BioRxiv.org).

### Data processing and Analysis

The data was trimmed to the interval of 21:00h to 05:00h to avoid shorter recordings due to battery failure taking place in some of the nights. However, flight bouts beginning before 05:00h and continuing after that, were included in full.

To characterize flight aptitude of individual mosquitoes we computed their total-flight duration between 21:00h and 05:00h (sum of all flight bouts per mosquito), longest flight-bout duration, and the total number of flight bouts per mosquito. Assuming that long-distance flyers would exhibit much higher values, compared with the majority of the population in at least one of these flight measures, we used median-based robust statistics, often used to detect outliers based on the distance of a value from the median in units of the median absolute deviation from the median (MAD). Unlike the mean and the standard deviation, the median and the median absolute deviation are not sensitive to extreme outliers and thus are considered “robust statistics” which can better represent the population in the absence of the excessive influence of these extreme values. For the same reason, they are better suited to detect outliers (Iglewicz et al. 1993; Leys et al. 2013; Cousineau and Chartier 2010; Rousseeuw and Hubert 2011). Following conventional guidelines, the threshold for outlier detection were values > 3.5 MAD units from the median were considered as “High Flight Activity” (HFA), as opposed to “Low Flight Activity” (LFA) or short-range flight, which represented values < 3.5 MAD units. Unless a population is observed to be “on the move” as are migratory swarms of locusts (Kennedy, 1951) or monarch butterflies (Gibo and Pallett, 1979), it is generally assumed that at any moment the fraction of individuals that express migratory behavior is small (Dingle and Arora 1973; Taylor et al. 1992; Taylor, Nault, and Styer 1993; Taylor et al. 2010; Lee and Leskey 2015; Menz et al. 2019). This assumption relies on Mark-Release-Recapture (MRR) studies suggesting that a sizeable proportion of the population remain near the area where they were marked, and also that the duration of the migratory phase last a few days and is typically shorter than the non-migratory phase, which can last several weeks or months (Johnson, 1969; Roff and Fairbairn, 2007). This view underlies our identification of mosquitoes exhibiting extremely high values of their flight aptitude measures as putative long-distance migrants.

## Results

A total of 707 individual mosquito recordings (including the manual extractions) from 114 assays were included in the analysis (Table 1). Individual flights longer than 10 minutes were recorded in 46 mosquitoes, while flights exceeding 30 minutes were recorded in 12 mosquitoes.

**Table 1.** Mosquito samples by season, gonotrophic state and species, for which flight data was collected and analyzed

| Season | Gonotrophic stage | *A. coluzzii* | *A. arabiensis* | *A. gambiae* | Total |
| --- | --- | --- | --- | --- | --- |
| Dec-Feb | Gravid | 62 | 0 | 2 | 64 |
|  | Unfed | 2 | 0 | 0 | 2 |
| Mar-Apr | Gravid | 31 | 0 | 0 | 31 |
|  | Unfed | 0 | 0 | 0 | 0 |
| Jul | Gravid | 75 | 0 | 3 | 78 |
|  | Unfed | 0 | 0 | 0 | 0 |
| Aug-Sep | Gravid | 125 | 10 | 62 | 197 |
|  | Unfed | 30 | 1 | 8 | 39 |
| Oct-Nov | Gravid | 114 | 95 | 76 | 285 |
|  | Unfed | 5 | 5 | 1 | 11 |
| Total |  | 444 | 111 | 152 | 707 |

Nightly flight activity and identification of putative Long-Distance Flyers (LDMs)

To determine if flight activity was concentrated in certain parts of the night, we examined hourly flight bouts, longest flight and total hourly flight (across species, season and gonotrophic state) (Fig. 1). To consider possible difference in nightly activity between appetitive and strong flyers, we evaluated both the median and 90^th^ percentile of each flight aptitude index.

**Figure 1.**
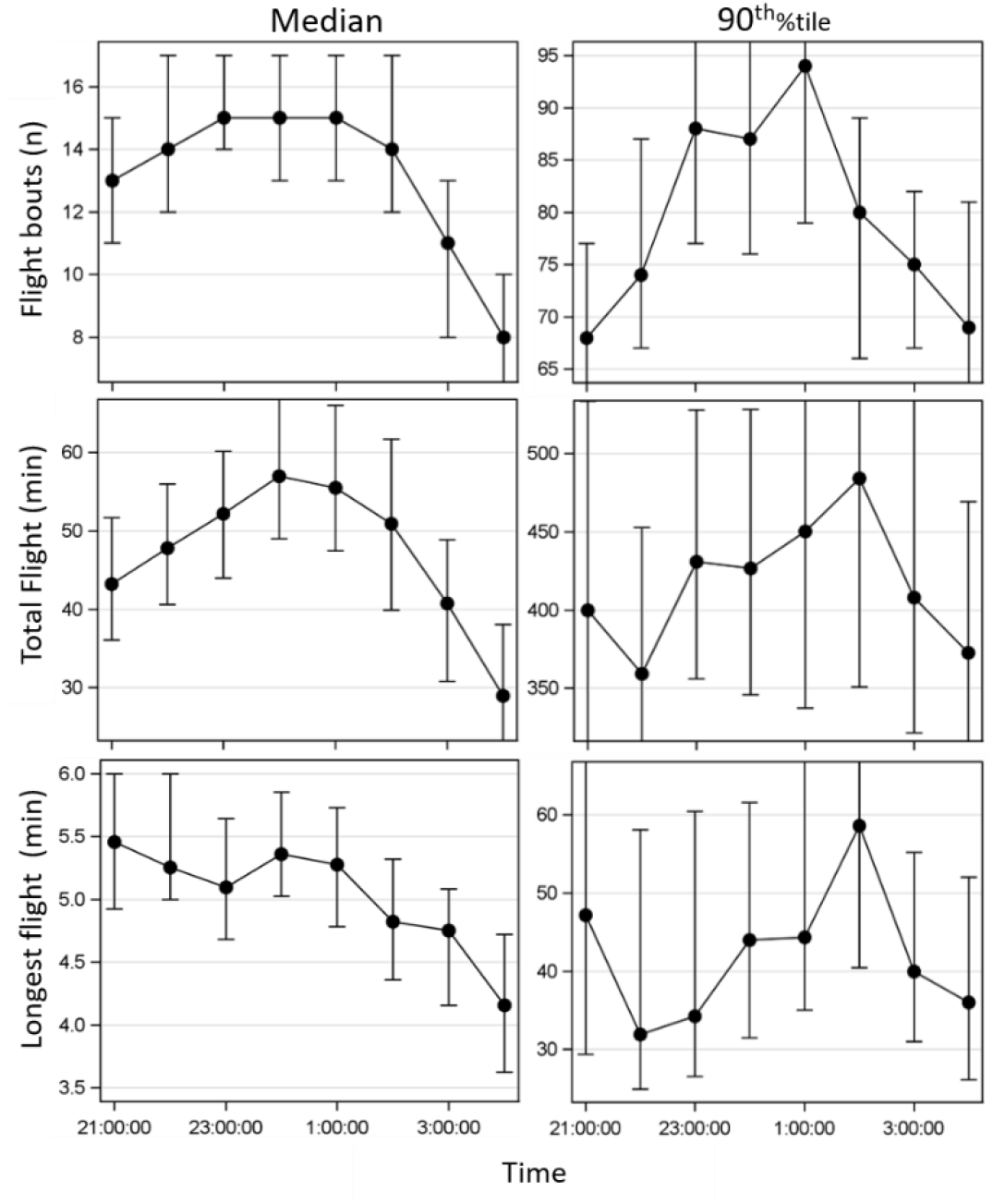
Nocturnal flight activity. Flight bouts and total hourly flight across all 707 mosquitoes, (across species, season, and gonotrophic state). Hourly flight activity (flight aptitude) through the night showing the median (left column) and the 90^th^ %il (right column), representing trends of most mosquitoes and higher flight activity mosquitoes respectively. The 95% confidence interval of each hour (based on bootstrapping) not shown in full to emphasize the nightly trend.

Overall, no significant peaks of activity were identified in the hourly flight data and variation, as measured by hourly 95% confidence intervals, was moderate (24-47% in median, and 25-45% in 90^th^ %il values). Although a mild trend suggestive of elevated total flight and flight bouts between 11:00 and 02:00h (but not in longest flight) was observed, it was not statistically supported as the 95% CI overlapped widely. We therefore concluded that the flight activity was spread rather homogenously across the assay time and used the full length/duration (21:00-05:00h) to measure flight aptitude.

### Identification of putative migrants

Considering all mosquito flights (n=707 mosquitoes), the distributions of each index of flight aptitude revealed high asymmetry having a long right-tail as is usual in frequency distributions of laboratory-measured bouts of flight (Davis, 1980; Johnson, 1976), with many individuals making short flights and a few making long ones. (Fig. 2).

**Figure 2.**
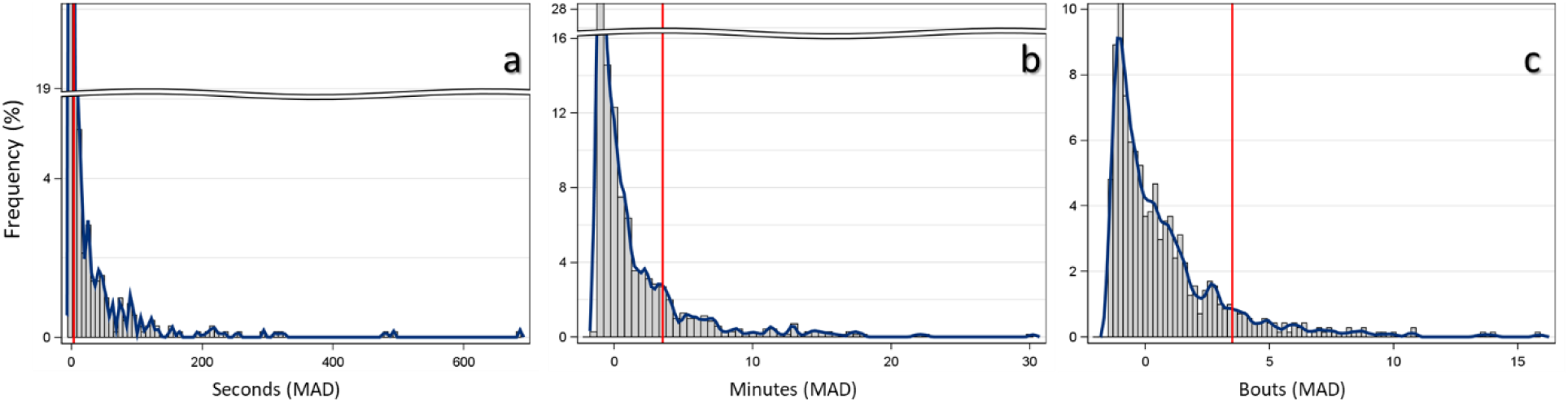
Flight aptitude index distributions: longest flight (a), total flight (b), and flight bouts (c). Flights were arbitrarily divided into two classes by units of deviation from the Median Absolute Deviation (x-axis; MAD units): LFAs; below 3.5 (left of vertical red line), and HFAs; above 3.5 (right of vertical red line). The data are based on 707 wild female mosquitoes representing three species, of both unfed and gravid females. Y-axis denotes the frequency in percent of the sample.

Following previous flight behavior studies (Dingle and Arora, 1973; Taylor et al. 1992; Reynolds and Frye, 2007; Lee and Leskey, 2015; Jones et al. 2016; Minter et al. 2018), the mosquitoes at the far-right end of the tail were suspected to represent long-distance flyers, or HFAs in our study. Based on conventional outlier detection algorithm (Iglewicz et al. 1993; Leys et al. 2013), an HFA mosquito was identified as presenting a flight aptitude value >3.5 MAD units of its median (see methods). In the following analysis, we evaluate differences in the proportion of HFAs among species, seasons, and gonotrophic state.

The flight aptitude indices were mostly significantly correlated with each other but correlation coefficients (Spearman) were negative (r=-0.43) between the longest flight and the number of flight bouts, or moderately positive (r=0.32—0.52) between total flight and the other indices (Fig. S3). The relatively low correlation coefficients indicate that each index contains unique information; in other words, flight bouts (describing “restlessness”) identifies a different type of HFA then those identified by the longest flight bout. Although both contribute to total flight and exhibit higher correlation coefficients with it (Fig. S3), we included all three indices as they capture, to a large extent, different aspects of HFA mosquitoes including restlessness and persistence of flight. At lease at the outset, it is difficult to determine which indices are more important predictors of migrants.

### Variation of flight aptitude by season

Seasonal variation in flight aptitude was tested on gravid *A. coluzzii*, which was the only species (and gonotrophic state) found across seasons. Significant differences were most pronounced in longest flight (P<0.002, overall Monte-Carlo Exact test) but were also detected in total flight (P<0.005, overall Monte-Carlo Exact test) (Fig. 3a-b respectively). In longest flight the highest fraction of HFA was discovered in the late wet season (Oct-Nov; 39.5%), followed by mid-wet season (Aug-Sep; 29%), early wet season (Jul; 19%) and early dry season (Dec-Feb; 22%), with the lowest being the late dry season (Apr; 10%, P<0.005, 2-tailed test, Fig. 3a). A similar trend, albeit with smaller differences, was detected in total flight with only a single significant difference between Oct-Nov and Apr (P<0.04, Fig. 3b). Flight bouts did not follow this pattern and showed no significant difference in the overall test (Fig. 3c).

**Figure 3.**
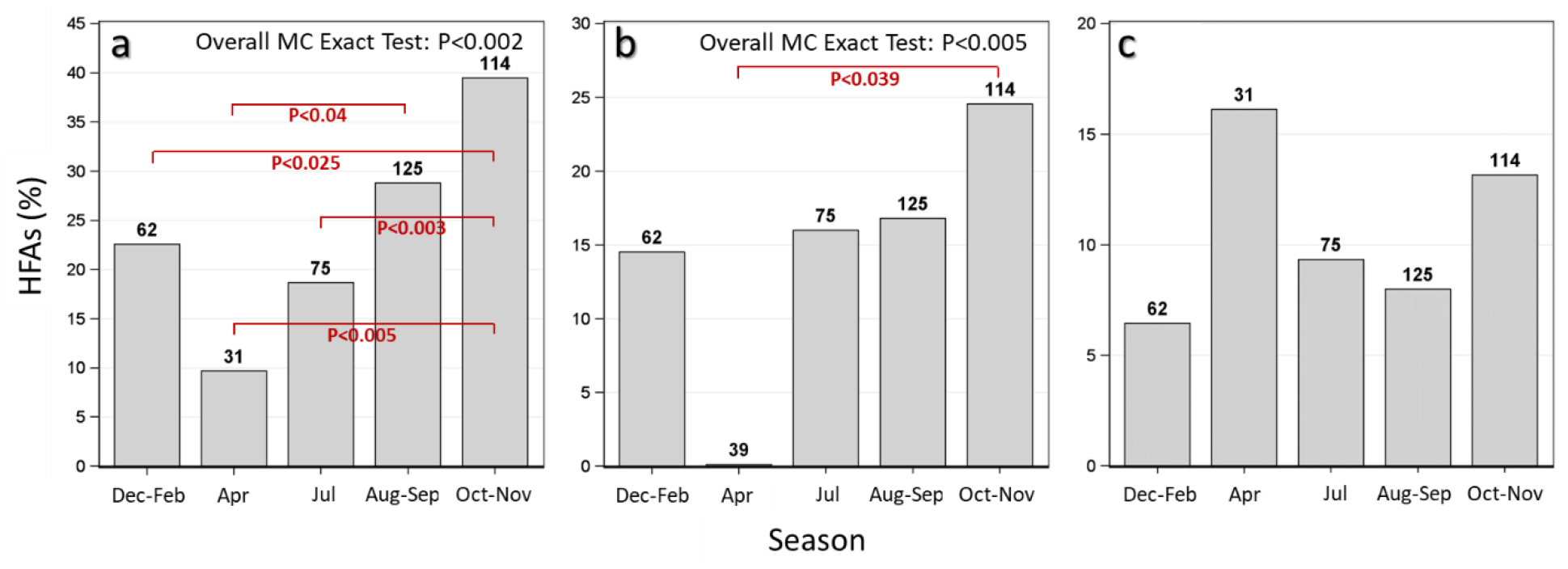
Variation of flight aptitude indices by season; longest flight (a), total flight (b), and flight bouts (c). Variation in *A. coluzzii* flight aptitude between seasons; The x-axis depicts the different parts of the year (‘seasons’); Dec-Feb and Apr represent the dry season. Jul, Aug-Sep and Oct-Nov represent the wet season. Y-axis values are percent frequencies of the HFA populations, with *n* above each bar. Seasonal flight aptitude comparison was carried out on gravid *A. coluzzii* females, the only species which had samples across seasons.

### Variation of FA between species

Variation in flight aptitude between species was tested on gravid females in Oct-Nov when all species were represented. Considering total flight, *A. coluzzii* exhibited a significantly higher fraction of HFAs (25%) than *A. arabiensis* (8%), with *A. gambiae* showing intermediate values (17%), (Overall Monte Carlo Exact Test χ^2^=10.0, P<0.015) (Fig. 4b). Contrasting test between the species showed a significant difference between *A. coluzzii* and *A. arabiensis* (Wald χ^2^=8.7, P<0.004). Although non-significant, similar trends were revealed in longest flight and flight bouts (Fig. 4a and c).

**Figure 4.**
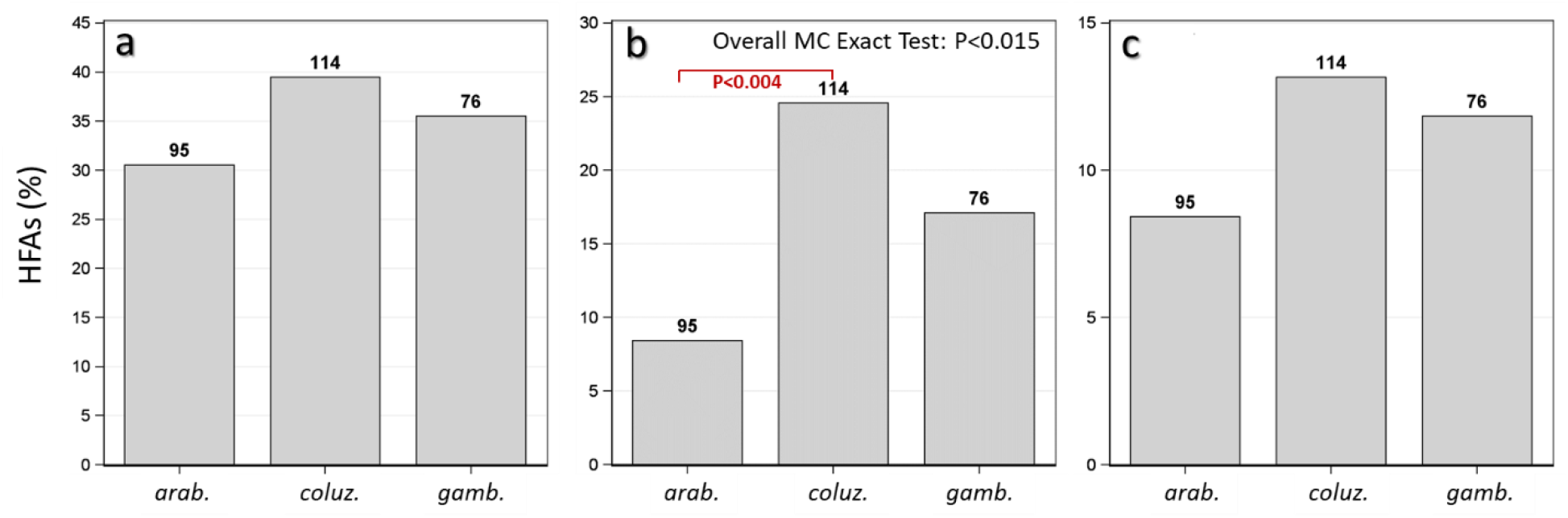
Variation of flight aptitude between species; longest flight (a), total flight (b), and flight bouts (c). Values are percent frequencies of the flyer populations, with *n* above each bar. Overall test is a contingency table exact test using Monte Carol with 10000 replicates. Unless otherwise specified all P-values pertain to 2-sided tests. Specific comparisons are shown were tested using contrasts in logistic regression if overall test was significant. Species comparison was done on gravid females in Oct-Nov, when sample size was suitable for comparison.

### Variation in flight aptitude between gonotrophic stages

The effect of gonotrophic stages on flight aptitude was tested after pooling the species and also with the stratification by species (CMH test) in August-September, when the number of unfed females was suitable for such a test. A significantly higher rate of HFAs among gravid females (11.2% vs. 0%, P<0.013 Overall Monte Carlo Exact Test) was detected in flight bouts (pooled; P<0.024, 1-tailed Fisher Exact test, and when stratified (CMH = P<0.049, 2-tailed test) (Fig. 5c). Although no significant differences were detected in longest- and total flight, they showed a consistent trend of higher HFA among gravid females (Fig. 5a and b respectively).

**Figure 5.**
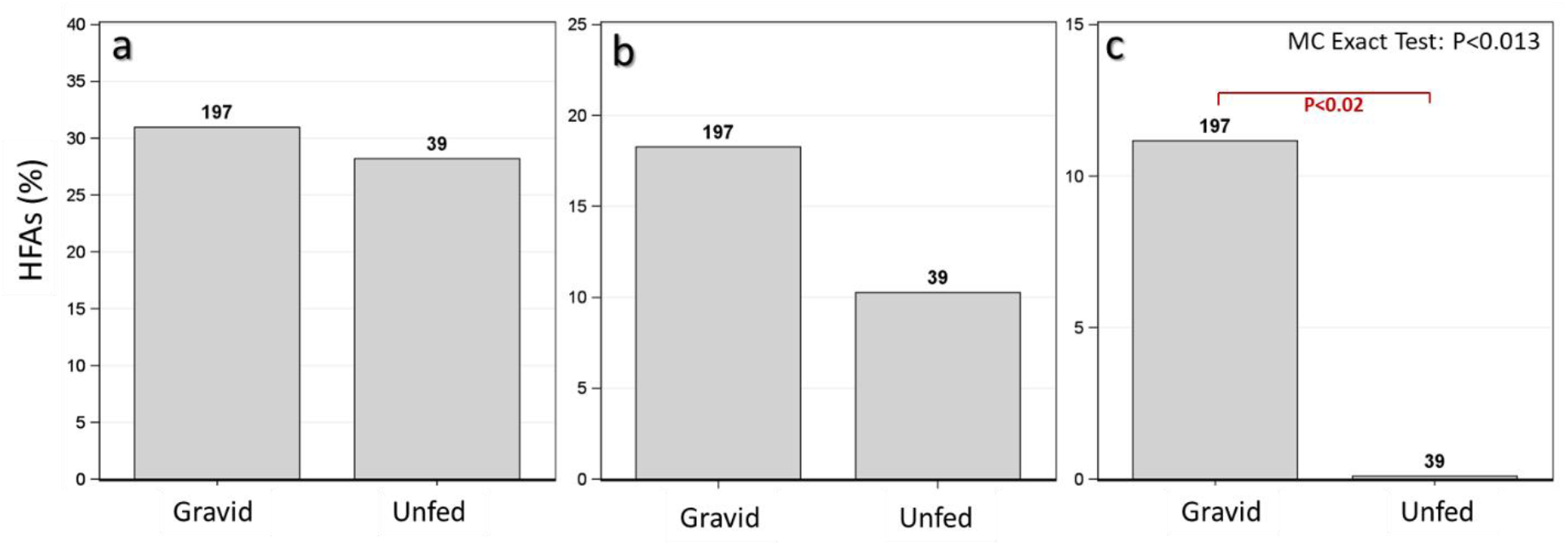
Variation in flight aptitude between gonotrophic stages; longest flight (a), total flight (b), and flight bouts (c). Overall test is a contingency table exact test using Monte Carlo with 10000 replicates. Unless otherwise specified all P-values pertains to 2-sided tests. This Gonotrophic state comparison was done when on pooled species in Aug-Sep, when sample size was suitable for comparison. Prior CMH test showed that the effect was significant across species (not shown) and no heterogeneity between species was detected.

### Wing morphology and flight aptitude

Significant differences were found in *A. coluzzii* in longest flight duration (Fig. 6a and d)and total flight (Fig. 6b and e), showing both longer (Fig. 6a and b) and wider wings (Fig. 6d and e) in HFAs (red) vs. LFAs (blue) (Overall ANOVA; P<0.02). Moreover, considering the hypothesis that HFAs have larger wings than LFAs, *A. coluzzii* HFAs, based on total flight had also significantly longer wings (P<0.043, 1-tailed test; Fig. 6b). Statistical significance of the same was higher using median and Wilcoxon tests comparing the medians of these groups (P<0.027, 1-tailed test) (Fig. 6b and e). Similar trends between HFAs and LFAs were seen in flight bouts albeit non-significant differences were found.

**Figure 6.**
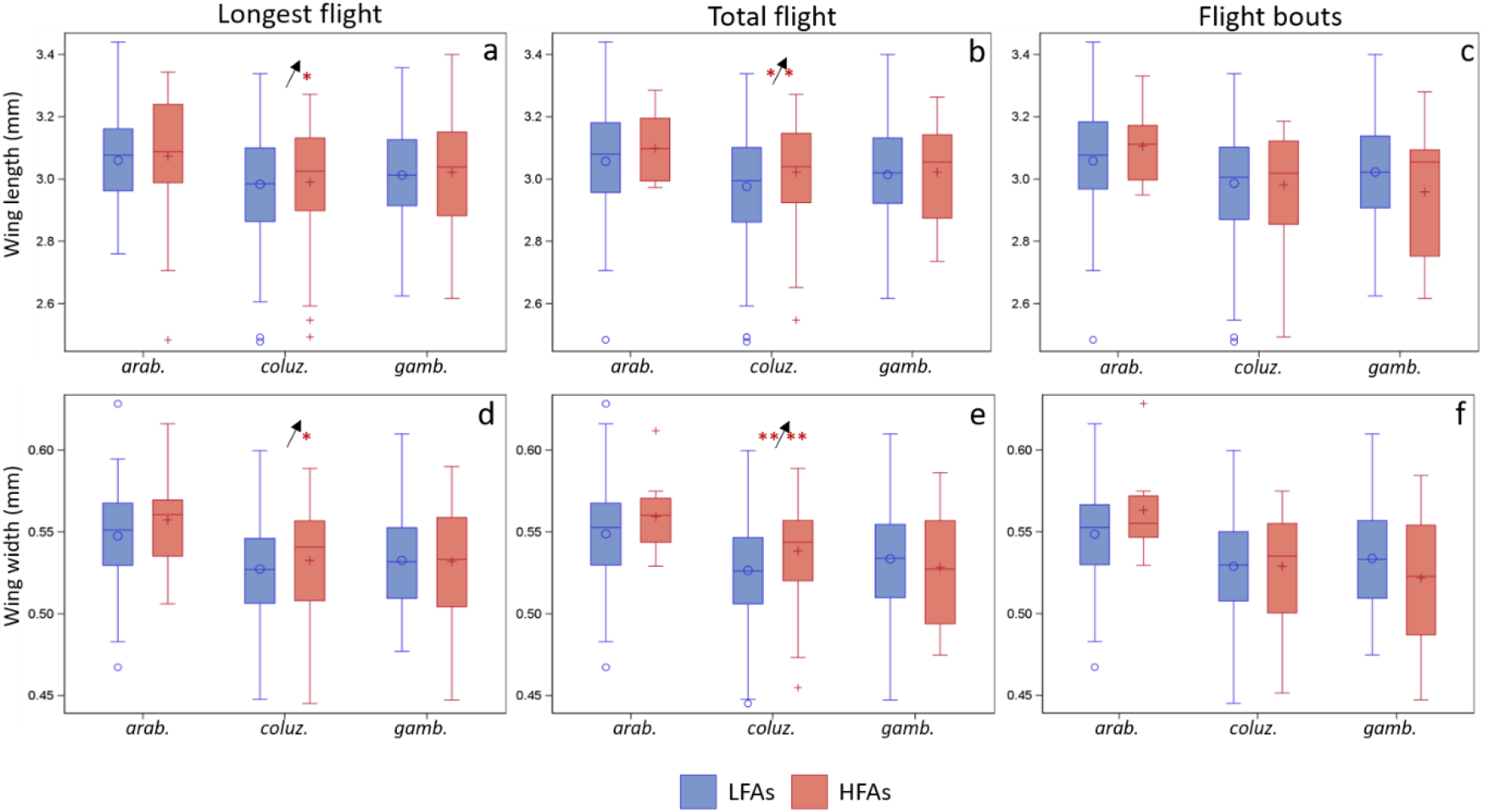
Box-whisker plot of wing length (top), and width (bottom) across the anopheline species (x-axis, abbreviated species) in longest flight (panels a and d), total flight (panels b and e) and flight bouts (panels c and f) for LFAs (blue) and HFAs (red). Mean marked as ○ (for LFAs) or + (for HFAs). Horizontal line within box is median. Box top and bottom are 25^th^ percentile and whiskers are upper and lower 75^th^ percentiles. *arab*. =A. arabiensis, coluz. = A. coluzzii., gamb. = *A. gambiae s.s.* Significantly larger mean and median wing dimensions in HFAs vs. LFAs indicated by asterisks on the left and right of the arrow (showing direction of increase), respectively. One tailed significance levels of P<0.05 and P<0.01 measured by ANOVA (left of arrow) or Median score tests (right or arrow); shown as ‘*’ and ‘**’ respectively.

### Allometry of wing to detect wing shape variation within HFA’s

In gravid *A. coluzzii* during the wet season, total-flight HFAs had wider wings than LFAs after accommodating wing length (P<0.035, 1-tail test). We found no significant interaction between wing length and HFAs, indicating that the shape effect was monotonic with wing length (Fig. 7). A similar trend was observed in *A. arabiensis*, but no significant difference in intercepts was detected, possibly due to smaller sample size.

**Figure 7.**
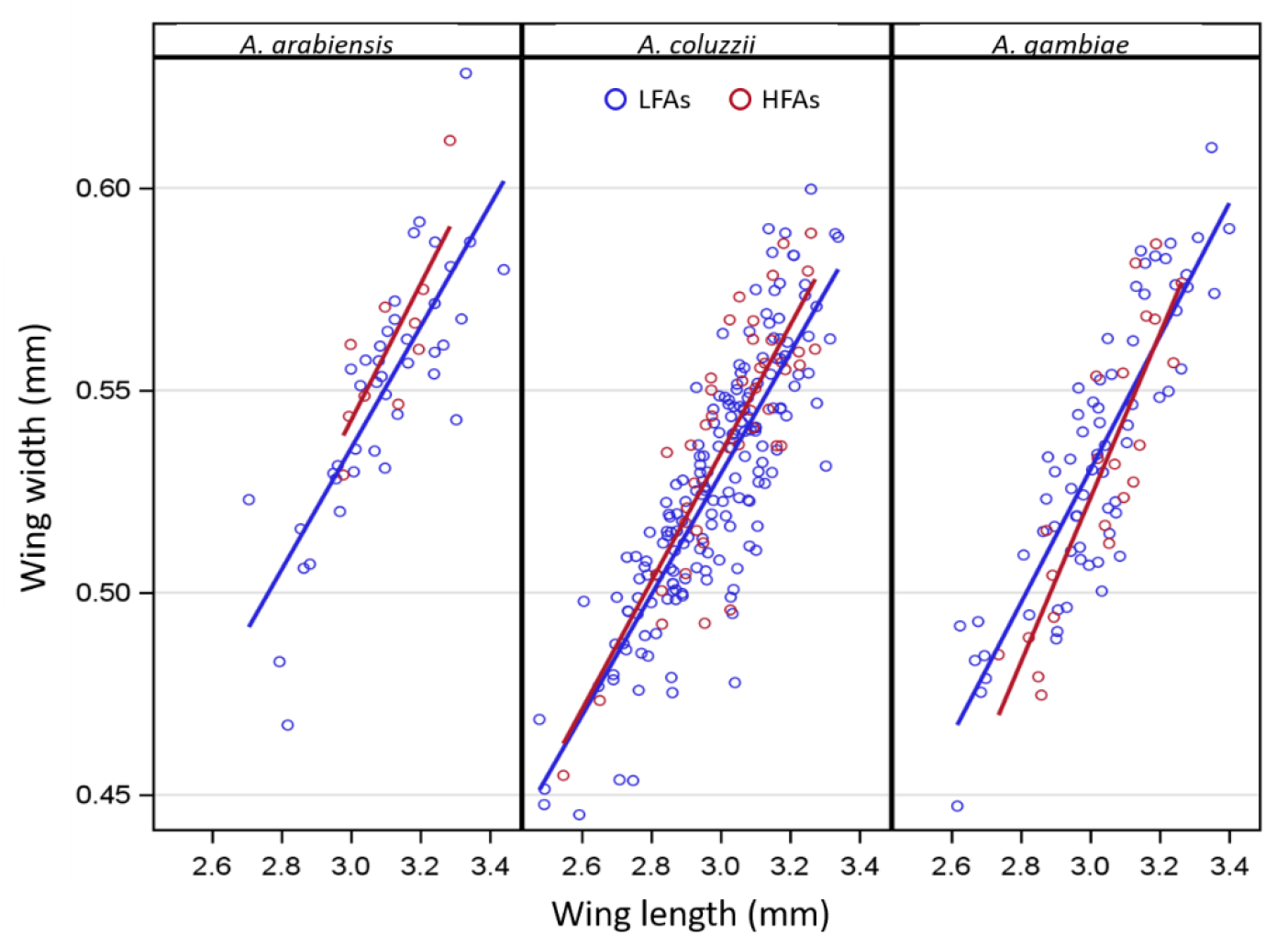
Wing allometry in HFAs (red) and LFAs (blue) for total flight, in all three species. In gravid *A. coluzzii* during the wet season, total-flight HFAs had wider wings than LFAs after accommodating wing length (P<0.035, 1-tail test).

Consistent with previous studies (Huestis et al. 2012; Arcaz et al. 2016), *A. coluzzii* showed longer wings in the early dry season (December-February) compared with later in the year (Fig. S4). This change was isometric, in other words, it was accompanied by a proportional reduction in wing-width during this time.

## Discussion

Recent studies on windborne migration of African malaria vectors raise new questions about this previously unrecognized behavior, including what fraction of the population engage in migration and how this fraction changes amongst species, across seasons, or ecological zones, with *Plasmodium* infection, etc. Identification of migrants in field experiments could help address such questions. In this paper, we subjected wild mosquitoes to a novel, field-adapted tethered-flight assay, to separate them into mosquitoes with high flight activity (HFAs) and low flight activity (LFAs). Following the approach used extensively in flight-mill studies, we presume that HFA is likely to be enhanced in long-distance migrants. Employing three flight aptitude indices measuring flight persistence (longest flight duration, and to a lesser degree, total flight) and restlessness (flight bouts), we evaluated the variation in the proportion of migrants in Sahelian populations of *A. coluzzii, A. gambiae* s.s., and *A. arabiensis*, over seasons and gonotrophic state. Although differences between groups were moderate, consistent with our predictions we found elevated HFA mosquitoes in the wet season and among gravid females. A prediction about species variation during the wet season was less certain. Based on Dao et al. (2014), we initially predicted migrants only in *A. gambiae* s.s. and *A. arabiensis*, however, based on the aerial sampling of Huestis et al. (2019), we predicted the presence of HFAs across all three species with higher HFAs in *A. coluzzii*, followed by *A. gambiae* s.s.. Consistent with Huestis and colleagues (2019), our results showed the highest HFAs in *A. coluzzii* and the lowest in *A. arabiensis*. During the wet-season, *A. coluzzii* HFAs exhibited larger wings than LFAs. Additional analysis indicated that wings of wet-season, *A. coluzzii* HFAs exhibited allometric change. In large part, these results agree with recent work, which showed the dominance of gravid *A. coluzzii* flying during the wet season at altitudes of 40-290 m above ground (Huestis et al. 2019).

Although an ultimate ‘comprehensive flight index’ and cutoff values to distinguish between long-distance migrants and appetitive flyers have yet to be found, ad hoc indices and values have been successfully used (e.g. Jones et al. 2016; Dällenbach, Glauser, et al. 2018; Minter et al. 2018). Flight mill-based studies to identify long-distance migrant insects often relied on total flight, longest flight and the number of flight bouts, and computed indices derived from those (Jones et al. 2016; Dällenbach, Glauser, et al. 2018; Minter et al. 2018; Naranjo 2019). We have selected those indices because of their demonstrated utility, intuitive appeal (it is difficult to conceive a long-distance migrant with low total flight), and because they portray distinct putative dimensions of migratory behavior: restlessness expressed by flight bouts, and flight persistence, expressed by the longest flight duration and by total flight duration. The low absolute value of the correlation coefficient between longest flight and flight bouts (r = −0.43, Fig.S3, bottom-right panel) highlights the independence of these indices. Only 10.5% of HFA mosquitoes based on longest flight were also classified as such by flight bouts (unlike longest- and total flight sharing over 90% of HFAs), again suggesting these modalities of flights are separate. Overall in our analysis, out of six comparisons (Table S2), HFA mosquitoes based on total flight revealed significant differences in five tests, followed by longest flight (three) and flight bouts (one), suggesting that persistence of flight is a more relevant modality for long-distance migration, similar to total duration, number of flights and longest flight duration (Jones et al. 2016). Notably, longest flight showed consistent trends with total flights in five (of six) comparisons, whereas flight bouts showed consistent trends in only two comparisons (Table S2).

### Flight aptitude variation over seasons, gonotrophic states and species

Previous work in the Sahel has shown that *A. coluzzii* populations build up rapidly after the first rains, i.e. May-June, and decline towards the late wet season (October), presumably entering aestivation (Dao et al. 2014). In contrast, both *A. gambiae* s.s and *A. arabiensis* populations that are altogether absent during the dry season, build up some 6 weeks after *A. coluzzii*, then vanish with the drying-up of surface water. Their population dynamics pattern suggests immigration from southerly sources where breeding sites are perennial. Based on these findings, we predicted elevated HFAs in both migratory species mostly during the wet season but minimal HFAs in *A. coluzzii*. Additionally, because long-distance migration in most insect species occurs before reproduction (Johnson, 1969; Dingle, 1974; Colvin and Gatehouse, 1993; Dingle and Drake, 2007b), we expected elevated HFAs in non-bloodfed as opposed to gravid females. However, recent results revealed that during the wet season in the Sahel, *A. coluzzii* predominated in high altitude aerial samples (n=23) compared with one *A. gambiae* s.s, and no *A. arabiensis* (Huestis et al. 2019), and that most of the mosquitoes were gravid. Accordingly, our revised expectation is that HFA will be higher in the wet season and in gravid mosquitoes. We now expected HFAs to be also present in *A. coluzzii*, but we could not form a unambiguous prediction about the differences among these species (Huestis et al. 2019). Overall, the results tend to agree with our predictions considering (*i*) elevated HFA during the wet season, (*ii*) elevated HFAs among gravid mosquitoes, and (*iii*) the presence of HFAs among all species, with highest flight aptitude in *A. coluzzii* and lowest in *A. arabiensis*. Uncertainty remains because of partial consistency concerning the different flight aptitude indices and coarse discrimination due to low statistical power among groups (Table S2). For example, the non-significant differences in *A. coluzzii* between the early dry season (Dec-Feb) and the late dry season (Mar-Apr, Fig. 3) or the non-significant difference between *A. gambiae* s.s. and the other two species. (Fig. 4).

### Wing morphometry and flight aptitude

Morphological differences between wings of HFAs and LFAs can provide strong evidence in support of the classification as well as indicate a distinct developmental program(s) for long-distance migrants. Our results reveal that during the wet season, *A. coluzzii* HFAs had a larger wing area due to an increase in wing width and length compared with the LFAs. The median values were correlated with the means but provided a higher degree of significance (P<0.027, 1-tailed test). These results were not confounded by variation in body size between seasons as the analysis was confined to the wet season, when no change in wing length was detected (Fig. S4). The larger wings of HFAs may reflect isometric increase in all aspects of body size or it may indicate an allometric change, e.g. an increase in wing area independent of body size. This is difficult to resolve with wild, mostly gravid mosquitoes because variation in dry weight may confound blood-meal size (and number) with body size. Indeed, during the early dry season (Dec-Feb), wings of *A. coluzzii* were inexplicably larger than other parts of the year (Fig. S4), consistent with previous results from two independent studies (Arcaz et al. 2016; Huestis et al. 2012). However, the allometric increase in wing width (over that expected by wing length) of *A. coluzzii* HFAs during the wet season further supports the validity of the classification, suggesting migrants may undergo a distinct developmental plan prior to adult formation.

### Interpretation of flight behavior

Members of the *A. gambiae* complex are possibly an example of ‘partial migrators’ (Dällenbach, Menz, et al. 2018, 2019), in which the majority of individuals do not engage in long-distance migration even during times when migration peaks. Moreover, after arrival, immigrants will exhibit LFA, therefore, the assay may identify only emigrants prior to their journey. Since the migratory phase may last only 1-3 days, we would expect to have only 1-2 days at most to capture a migrant before she embarks on her journey. Thus, a large sample size is required to represent migrants among the more numerous ‘appetitive flyers’ which add to noise in the data and limit the statistical power of detecting differences.

No information is available to date concerning mosquito flight behavior at high altitudes. Even if tethered flight assays accurately identify long-distance migrants, the flight data generated in the assay is unlikely to mirror exactly the flight behavior at altitude. For example, the total flight duration may greatly underestimate actual flight in altitude – simulations based on the aerial sampling data (Huestis et al. 2019) suggest night-long migratory flights in some cases. Likewise, flight mill results may fail to exhibit flight patterns matching expectations based on migration because of technical as well as biological reasons (Taylor et al. 2010; Jones et al. 2016; Minter et al. 2018; Naranjo 2019). Fair examples are lack of lift generation and lack of tarsal contact which may lead to unrealistically extended flight while, on the other hand, the lack of sensory cues from air movement or apparent ground movement may curtail flights. Our flight aptitude assay relied on fixed-tethered mosquitoes placed in tubes to partially isolate them from surrounding environmental cues. Thus they may express intrinsically-driven flight as previously suggested in works on locusts and moths (Bazazi et al. 2012; Reynolds et al. 2015). Finally, flight assays for migrant identification will be more informative when additional information is gathered on each mosquito to assess agreement with other aspects of the migration syndrome (Lin et al. 2016; Jones et al. 2015). For example, combining data on nutritional reserves (typically elevated before migration), response to host or oviposition site cues (typically inhibited prior and during migration) (Slansky, 1982), cuticular hydrocarbons (presumably elevated prior to migration to enhance desiccation tolerance) (Arcaz et al. 2016), morphometrical analysis of size and shape of wings, thorax, and spiracles, and transcriptome analysis.

## Declarations

### Consent for publication

All authors have read and approved the final version and consent to its publication.

### Availability of data and materials

The datasets during and/or analysed during the current study available from the corresponding author on reasonable request.

### Authors’ contributions

Conception and design of the work: RF, ASY, MD, AD, SD, ZLS and TL. Acquisition of data: ASY, MD, AD, SD, MS and ZLS. Analysis and interpretation of data: RF, BJK and TL. Drafted the work: RF and TL. Substantively revised the work: RF, TL, ASY, MD, AD, SD, ZLS, MS and BJK.

### Ethics approval and consent to participate

Not Applicable.

## Acknowledgements

We thank the people of the villages of Thierola and Selingue in Mali for their support of the field work. We wish to thank Drs. Don R. Reynolds, Jason W. Chapman and Christopher M. Jones for valuable discussions and comments on previous versions of this manuscript. We thank Drs. Manu Prakash, Felix Hol, Haripriya V. Narayanan and Deepak Krishnamurthy for valuable discussions on flight mechanics. We wish to thank Samuel Moretz, André Laughinghouse, Kevin Lee, Anish Prasanna for providing logistical support. Thanks to Dr. Xavier Martini for providing a flight mill for testing. This study was supported by the Division of Intramural Research, National Institute of Allergy and Infectious Diseases, National Institutes of Health, Bethesda MD.

