## Supplemental data for "Quantifying flight aptitude variation in wild *A. gambiae* s.l. in order to identify long-distance migrants"

Supplementary information file

Wing measurements

For each wing 14 specific landmarks (Fig. S1), i.e. longitudinal vein intersections, with either the wing edge or between themselves, were mapped using the tps-DIG 2.15 software package (Rohlf, 2010). Wing length (measured between points 1 and 10) and wing width (see below) were used as the primary measure of body size and wing area throughout this analysis. Wing width was derived from the average value of the height of three triangles formed between landmarks: 1-2-13 (proximal triangle, green), 3-12-13 (medial triangle, black), and 2-5-11 (distal triangle, blue) (Fig. S1).

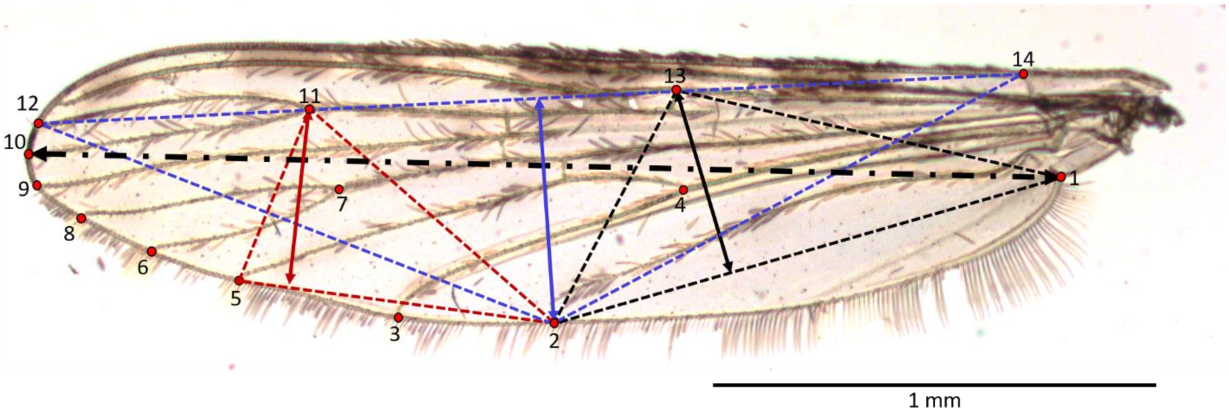


Figure S1. Wing of *An. gambiae* s.l. (x25 magnification); Landmarks (14) denoted by numbers. Black dot-dash line represents the wing length between landmarks 1-10. Wing width was the average value of the height of three triangles formed between landmarks: 1-2-13 (proximal triangle, black), 2-5-11 (distal triangle, red), and 2-12-14 (medial triangle, blue). Scale bar = 1 mm.

Tethered flight assay

Using entomological pins (Morpho No.3. Ento Sphinx, Czech Republic), sharp ends were clipped off, pins were bent twice at 90°(to a square bracket shape). The tip of one end of the pin was lightly dipped in white glue (Elmer's, Glue-All E1322, Atlanta, GA) and gently pressed on to the ventral side of the posterior abdomen. The other end of the pin was threaded through the base of a disposable 10 µl pipette tip of which the nozzle cone was cut off (Fig. S2a).

Tethered mosquitoes were allowed to fully recuperate from the ether for 1-2 hours before the flight assay (moving of legs and wings usually began 2-3 minutes after removal from the ether), during which time their fore-legs were allowed to rest on a folded piece of paper (='leg-rest') providing tarsal contact and preventing flight (Fig. S2b). At the end of the recuperation time, tethered mosquitoes were inserted into individual flight tubes (n=18, 50 ml Falcon ®, Corning, NY, USA) positioned within a polyurethane foam hive (cut-up mattress) for improved soundproofing and environmental cue reduction (Fig. S2c). Tether constructs (mosquito, pin and base) were secured onto a small piece of double-coated urethane foam tape (3M®, Cat. No. 4026. St. Paul, MN) to fasten them 1 cm inward of the flight tube edge. Each flight tube housed a small microphone (ME-15, Olympus America Inc., Center Valley, PA, USA) attached to a portable voice recorder (VN-5200PC, Olympus America Inc., Center Valley, PA, USA) (Fig. S2c*i*) to record flight sound.

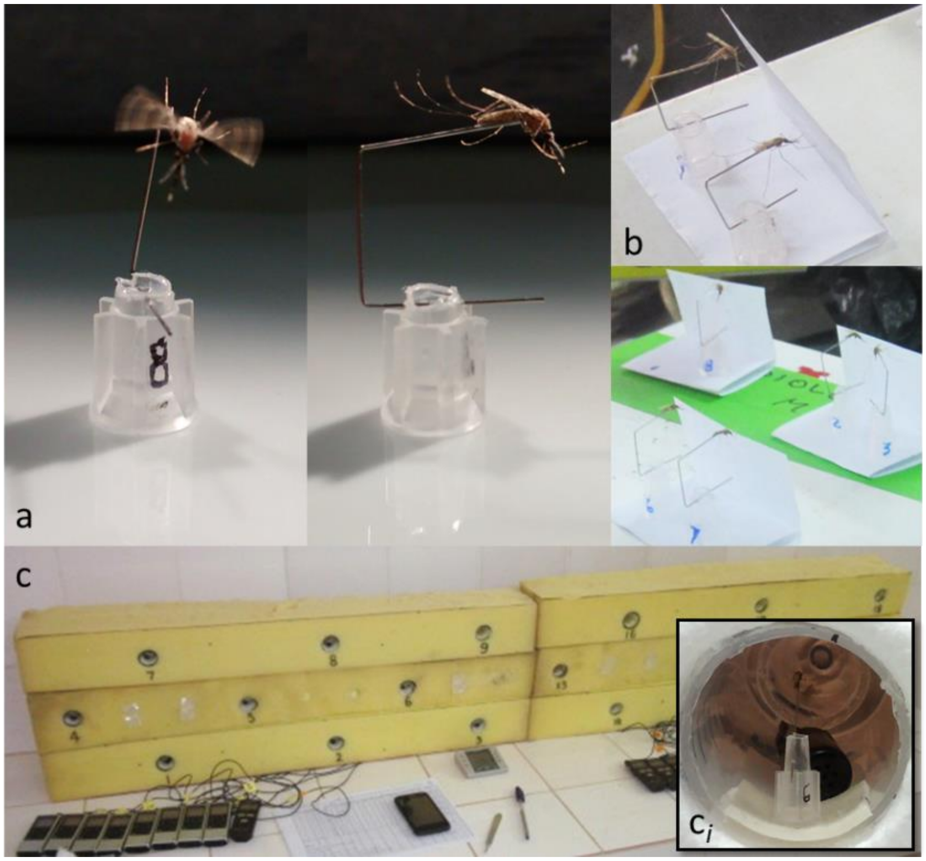


Figure S2. Tethered female *Anopheles gambiae* s.l. (a) Entomological pin attached to ventral side of posterior abdomen of the mosquito (a; right), allowing unobstructed flight (a; left). Bottom part of pin inserted into clipped 10 µl pipet tip as a base. (b) Tethered mosquitoes in recuperation time before assay start, fore-legs resting on folded paper for tarsal contact preventing flight. (c) Tether flight hive of 18 flight tubes housed inside soft (mattress) foam for surrounding sound muffling and external cue reduction. Each flight tube microphone connects to an individual sound recorder. ($c_i$) Tethered female inside flight tube (polystyrene prototype, not used in the experiment); tethered mosquito construct attached on to double-sided foam tape with microphone (black) in backdrop. Photos: a and $c_i$;RF, b and c; ASY.

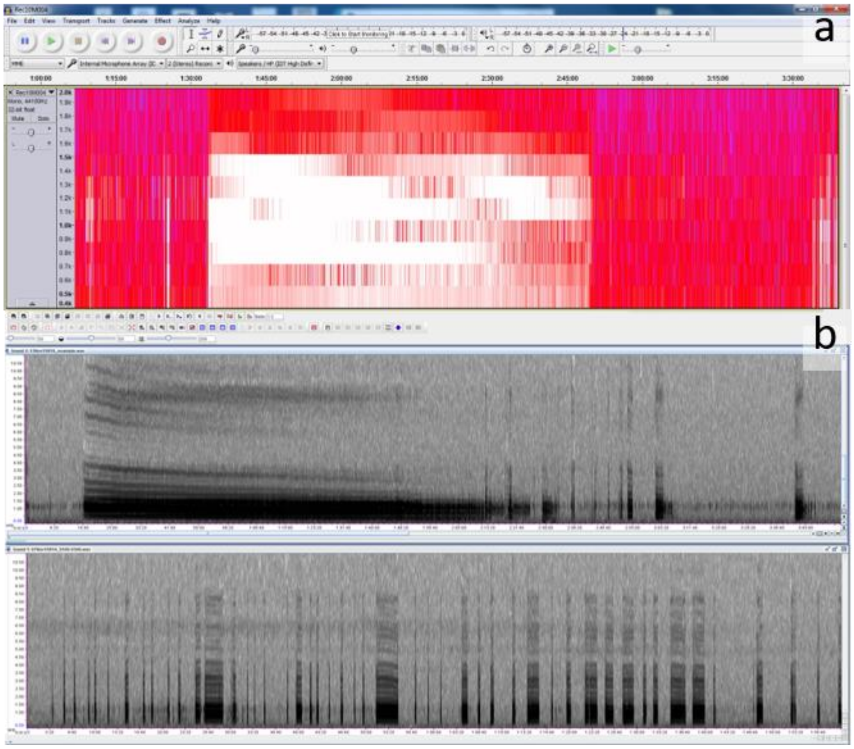


Figure S3. Flight sound file output (spectrogram view) in Audacity (a) and Raven Pro 1.5 software (b). Energy frequency (y-axis) and time of flight during the assay (x-axis) allowed extraction of flight durations during the assay. Panel a shows continuous flight bout of >70 minutes (in white), followed by a >40-minute rest period (in red). Panel b (top) shows >120-minute long continuous flight (in black) ending in several shorter bouts, whereas the bottom panel shows multiple bouts spanning 2-20 seconds.

### Flight Sound Extraction

Flight bouts were identified visually in spectrogram view (Fig. S2d) on Audacity (https://www.audacityteam.org/) or Raven Pro 1.4 software (ref), and questionable flights were confirmed by listening for flight sound directly on the sound file.

### Nocturnal flight distribution

Figure S3. Nocturnal flight (log scale) activity per species (color), season and FA index (column). Numbers on graphs indicate sample size per species.

Table S1. Summary data of all wild tethered mosquitoes in the final analysis. Number of mosquitoes by season, gonotrophic state and species.

| Season | Gonotrophic State | *A. coluzzii* | *A. gambiae* | *A. arabiensis* | Total |
|---|---|---|---|---|---|
| Dec - Mar | Gravid | 62 | 2 | 0 | 64 |
| | Unfed | 2 | 0 | 0 | 2 |
| Apr | Gravid | 31 | 0 | 0 | 31 |
| Jul | Gravid | 75 | 3 | 0 | 78 |
| Aug - Sep | Gravid | 125 | 62 | 10 | 197 |
| | Unfed | 30 | 8 | 1 | 39 |
| Oct - Nov | Gravid | 114 | 76 | 95 | 285 |
| | Unfed | 5 | 1 | 5 | 11 |
| Total | | 444 | 152 | 111 | 707 |

Flight Aptitude Indices relationships

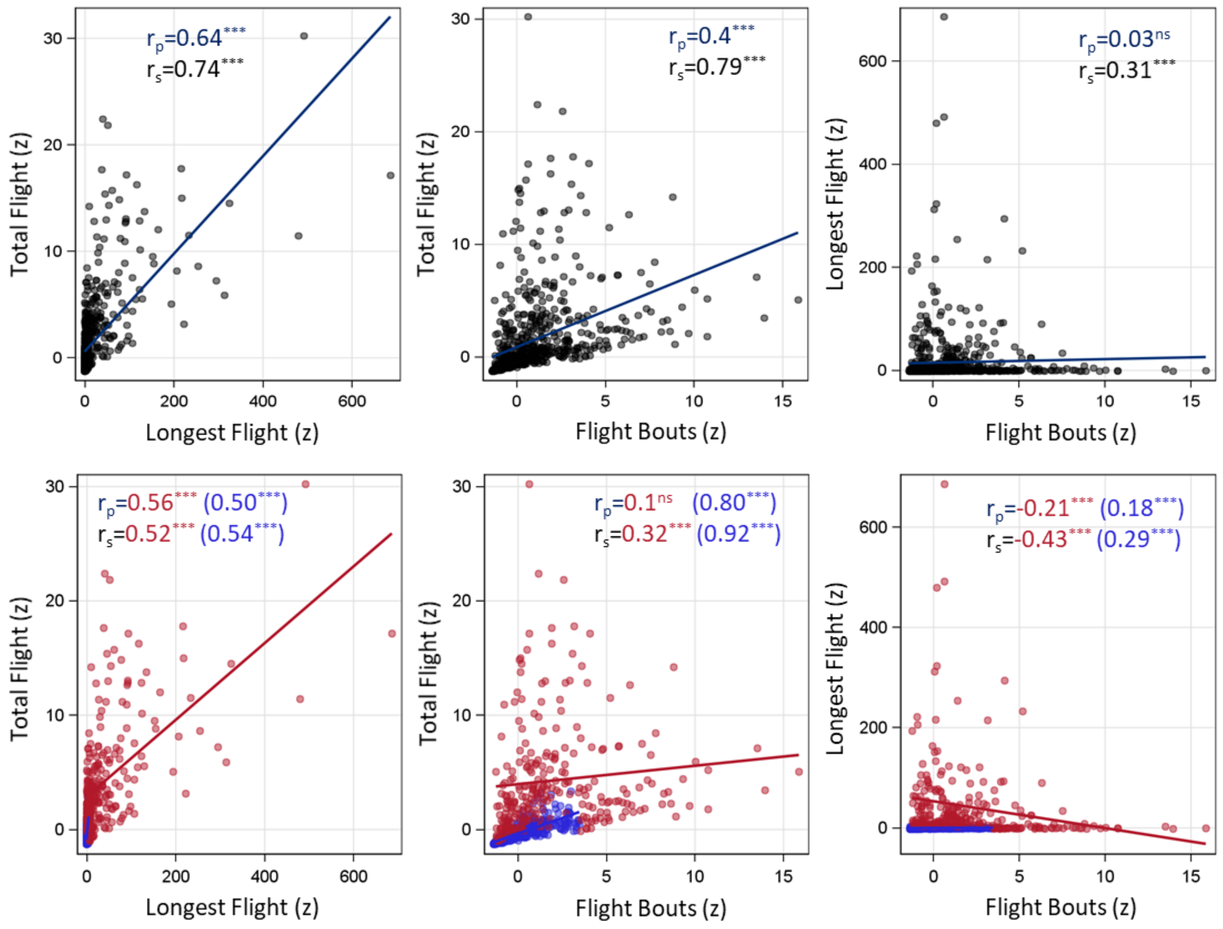


**Figure S3.** Spearman and Pearson correlations between the key flight aptitude indices: total flight, longest flight and flight bouts. Upper row: scatter plots of flight aptitude indices pairs showing linear regression as trend lines (dark blue) across all data (N=707 mosquitoes). Pearson and Spearman correlation coefficients are given with their significance level (*** and ns denotes P<0.001 and P>0.05, in Pearson (p) and Spearman (s) coefficients respectively).

Lower row: Scatter plots as above, showing the relationships for HFA mosquitoes (red; n=261) and LFA mosquitoes (blue; n=446). HFAs were defined based on any one significant flight aptitude index (or more).

Wing size and Flight Aptitude

Wing length was putatively selected between the two points on the wing best representing its longest aspect (points 1 and 10). The wing width on the other hand, has no agreed upon set land marks, and no two set points on the wing represent its width loyally. To overcome this, we selected several pairs of points on the wing, representing the proximal, medial and distal sections, and providing different distances. Assessing the degree of correlation between the three widths we eventually selected the mean of the three as the wing width.

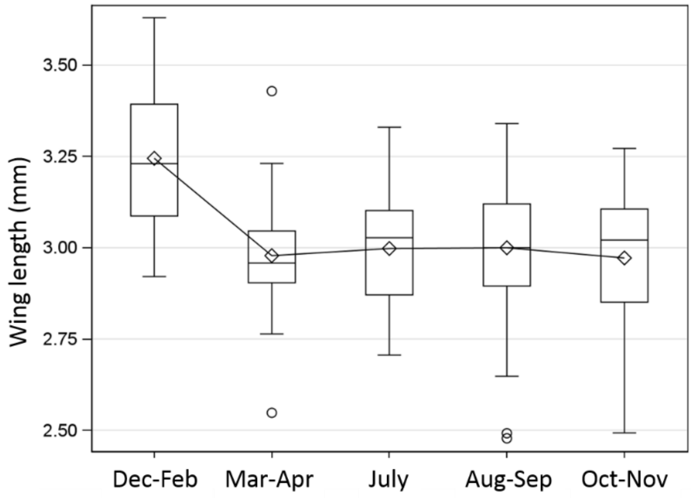


Figure S4. Wing length in *Anopheles coluzzii* by time of year (season). Mean length represented as diamond symbols (◊). Medians as horizontal lines inside boxes.

Table S2

| **Group tests** | **Total Flight** | **Longest Flight** | **Flight Bouts** |
|---|---|---|---|
| Season [a] | * | ** | reverse trend |
| Species [b] | * | similar trend | similar trend |
| Gonotrophic state [c] | similar trend | similar trend | * |
| Wing morphology (*A. coluzzii*, wet season) | | | |
| Wing length | ** | * | reverse trend |
| Wing width | ** | * | reverse trend |
| Allometry | * | reverse trend | reverse trend |
| HFA Shared Frequency (%) [d] | | | |
| Total Flight | 100 | - | - |
| Longest Flight | 90.3 | 100 | - |
| Flight Bouts | 23.9 | 10.5 | 100 |
| HFA Overall Frequency (%) [e] | 16.0 | 29.7 | 10.0 |

* significance of P<0.05, ** significance of P<0.001
[a]Oct-Nov > Aug-Sep = Jul = Dec-Feb > Mar-Apr
[b]A. coluzzii > A. gambiae > A. arabiensis
[c]Gravid > Unfed
[d] Frequency of HFA mosquitoes in two FA indices
[e] Total frequency of HFAs per FA index
